# Basal extrusion drives cell invasion and mechanical stripping of E-cadherin

**DOI:** 10.1101/463646

**Authors:** John Fadul, Gloria M. Slattum, Nadja M. Redd, Mauricio Franco Jin, Michael J. Redd, Stephan Daetwyler, Danielle Hedeen, Jan Huisken, Jody Rosenblatt

## Abstract

Metastasis is the predominant reason that patients succumb to cancer, yet the mechanisms that drive initial tumor cell invasion are poorly understood. We previously discovered that crowding-induced apical extrusion drives most epithelial cell death, critical to maintaining constant cell densities. Oncogenic mutations can disrupt apical cell extrusion, instead causing masses to form and aberrant basal extrusion. Using transparent zebrafish epidermis to model simple epithelia, we can image invasion events live at high resolution. We find that KRas/p53-transformed cells form masses and, at completely independent sites, invade by basal extrusion. Basal extrusion also causes invading cells to simultaneously mechanically shed their entire apical membranes and E-cadherin. Once cells invade the underlying tissue, they migrate throughout the body, divide, enter the bloodstream, and become different cell types. KRas-transformation makes cells intrinsically invasive by increasing basal extrusion rates; collaborating mutations in p53 allow disseminated cells to survive at distant sites.

## INTRODUCTION

Metastasis is tightly linked with poor cancer prognosis. For a cancer cell to metastasize, it must first invade from the epithelium, where ~90% of tumors originate. It has long been assumed that cancer cells use signaling and mechanisms that drive cells during development to undergo an epithelial to mesenchymal transition (EMT). While cancer cells may adopt several developmental features, they do not do not fully recapitulate normal differentiated lineages that contribute to healthy functional tissue but instead produce partially dedifferentiated malfunctioning cell types that disrupt organ function. Moreover, circulating tumor cells and metastatic cancer cells typically carry both mesenchymal and epithelial markers, rather than purely mesenchymal markers. The classical developmental EMT transcription factors, Twist, Snail, and Zeb, are dispensable for pancreatic cancer metastasis, further suggesting that cancers do not rely on developmental EMT programs to promote their spread (Zheng et al., 2015).

We previously discovered a process called cell extrusion that drives most epithelial cell death to maintain constant cell densities (Eisenhoffer et al., 2012; Eisenhoffer and Rosenblatt, 2013; Gudipaty et al., 2017; Rosenblatt et al., 2001). When cells within an epithelium become too crowded, crowding forces activate the stretch activated channel Piezo1 to produce the lipid Sphingosine 1-Phosphate (S1P), which signals the S1P2 receptor to extrude live cells out of the epithelium (Eisenhoffer et al., 2012; Gu et al., 2011; Slattum et al., 2009). After these cells extrude out apically and detach from the underlying matrix and its associated survival signaling, they die by anoikis, or apoptosis due solely to loss of survival signaling. In this way, crowding-induced extrusion simultaneously prevents gaps from forming in an epithelial sheet while keeping cell numbers constant within an epithelium. Cell extrusion is critical for maintaining a normal, healthy epithelial cell density, since we found that oncogenic mutations in either Adenomatous Polyposis Coli (APC) or KRas (Marshall et al., 2011; Slattum et al., 2014) or loss of S1P_2_ receptor (Gu et al., 2015) that are critical drivers of pancreatic, lung, and colon cancers disrupt apical extrusion signaling. In all cases where apical extrusion is disrupted, cells accumulate into chemotherapy-resistant masses or extrude aberrantly, basally, under the epithelium, instead of extruding out apically to die (Gu et al., 2015). If cells extrude apically, they will be eliminated through the lumen even if they do not die; if they extrude basally, they could be retained within the body and take on another fate. For example, during Drosophila development, neuroblasts basally extrude or delaminate, enabling them to de-differentiate and re-differentiate into neurons (Hartenstein et al., 1994; Simoes et al., 2017). Because oncogenic mutations that disrupt apical extrusion drive aggressive cancers prone to metastasis (Fadul and Rosenblatt, 2018; Gudipaty and Rosenblatt, 2017; Slattum and Rosenblatt, 2014) and rescue of S1P_2_ dramatically reduces both tumors and their metastases in a mouse orthotopic pancreatic cancer model (Gu et al., 2015), we wondered if basal extrusion could initiate tumor invasion.

To test if aberrant, basal extrusion could cause tumor cell invasion in vivo, we mosaically expressed a fluorescently-tagged oncogenic, activating mutation in KRas^V12^ that disrupts apical extrusion signaling in zebrafish embryonic epidermis. Currently, most cancer studies use mice, which offer excellent genetic models that recapitulate many aspects of human disease. However, mouse models require intravital imaging, which introduce wounds that could affect cell behavior, and preclude our ability to visualize cells stochastically invading from epithelia. Zebrafish epidermis models the simple epithelia where carcinomas originate and circumvents the problems associated with mouse studies, allowing us to film all cells live at sub-cellular resolution (Eisenhoffer and Rosenblatt, 2011; Eisenhoffer et al., 2017). We find that cells expressing GFP-KRas^V12^ in the outer epidermal layer form masses at sites where wild-type cells normally extrude apically to die and, at completely separate sites, aberrantly basally extrude into the underlying tissue. Basal extrusion not only drives a new mechanism for tumor cell invasion but also simultaneously sloughs of the entire apical cell surface, including E-cadherin, a critical epithelial determinant. Following invasion and loss of epithelial identity, basally extruded cells can migrate throughout the zebrafish body, divide, and become different cell types. Although KRas mutations are generally considered to be early mutations in aggressive cancer types, our findings suggest that they impart tumor cells with an intrinsic metastatic propensity, but that they will only survive dissemination if they can escape cell death. Moreover, invasion by basal extrusion occurs at sites distinct from the primary masses, which could impact the way that we diagnose and treat these aggressive cancer types.

## RESULTS

### Epidermal KRas^V12^ cells form masses, basally extrude, and invade

We tested the fate of epidermal cells mosaically-expressing EGFP-KRas^V12^ or EGFP-CAAX, both directly tethered to the membrane by farnesylation (Schmick et al., 2014), by injecting DNA plasmids driven by the krt4 promoter into one-cell stage AB wildtype embryos. KRas^V12^ expression caused masses to form at the zebrafish fin edges where cells typically extrude out to die as they converge and become crowded (Fig. 1 A, B, & D). Each tumor-like mass was comprised of between 3-40 cells. While KRas^V12^-expression caused 58.0% (58 of 100 embryos) of embryos to form cell masses, only 1% of CAAX-expression did (only 3 small masses of 290 embryos), despite being routinely more widely expressed than EGFP-KRas^V12^ (Fig. 1 A, C).

**Figure 1.**
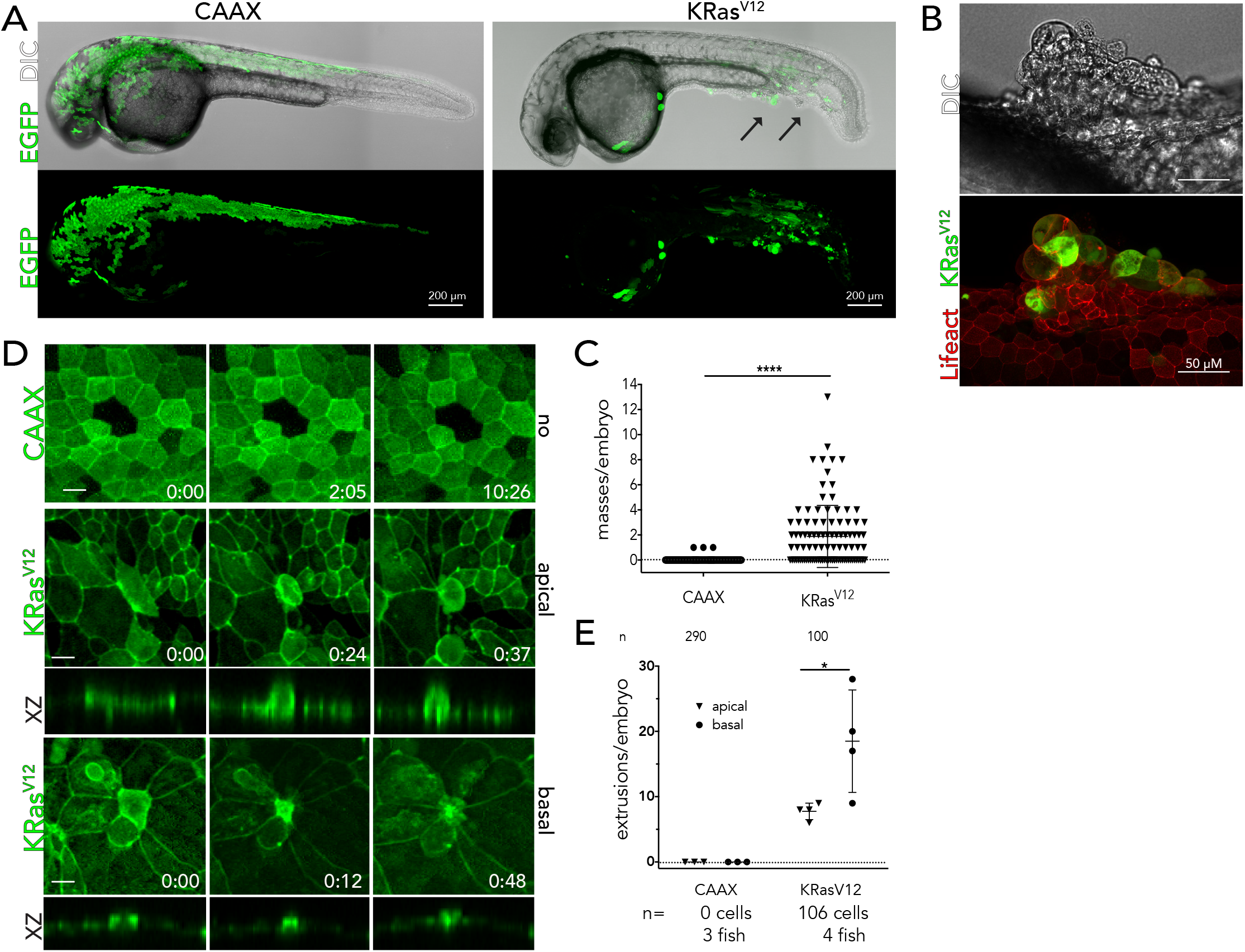
KRas^V12^ induces formation of epidermal cell masses and basal extrusion in wildtype zebrafish embryos. (A) 26 hpf AB wildtype zebrafish embryos injected with krt4-driven EGFP-CAAX or KRas^V12^ at one-cell stage. Epidermal cell masses are indicated by black arrows. (B) Large mass of EGFP-KRas^V12^ cells. (C) Quantification of cell masses per embryo. (D) Time-lapse imaging of krt4-driven EGFP-Lifeact (green) with mosaic expression of EGFP-CAAX where cells do not change over time (from SMovie1), or of EGFP-KRas^V12^ where cells extrude apically (from SMovie 2) or basally (from SMovie 3), with XZ sections beneath showing that the constricting F-actin on the basal or apical surface of the extruding cell, respectively (hh:mm and scale bars = 10μm). (E) Quantification of apical and basal extrusions in EGFP-CAAX or KRas^V12^ cells. For all graphs, values are from the Ns listed below the graph, where error bars = s.e.m. P-values from unpaired T-tests compared to control are *^****^<0.0001, ^*^<0.01.*

Additionally, many of the KRas^V12^ cells that did not form masses were significantly more likely to extrude than control cells, potentially accounting for the reduced number of KRas^V12^ cells over time. When EGFP-KRas^V12^ cells extruded, they predominantly extruded basally back into the body of the zebrafish (Fig. 1 D&E). Expression of KRas^V12^ in zebrafish lines containing krt4:Lifeact:mCherry or periderm:Lifeact-EGFP (Eisenhoffer et al., 2017) enabled us to visualize the intercellular actomyosin ring formed between the extruding and neighboring cells (Rosenblatt et al. 2001, Slattum et al. 2009) (Fig. 1D). Analysis of cell extrusions in movies of zebrafish between 24-46 hpf confirms that while no CAAX cells extrude, KRas^V12^ cells extrude at high rates (Fig. 1 D, E and associated SMovies 1-3). While apically extruded cells are essentially eliminated from the zebrafish, we hypothesized that basally extruded KRas^V12^ cells may be able to survive and invade the underlying matrix and tissue.

Using confocal imaging of fixed, immunostained zebrafish, we found numerous EGFP-KRas^V12^ cells inside the bodies of zebrafish at 48 hpf, whereas EGFP-CAAX cells remain at the epidermis (Fig. 2 A-C). We scored invaded cells as any EGFP-positive cells found beneath the basal layer of epidermal cells, which were labeled with p63 (Fig. 2 A-C, Smovie 4). We excluded from analysis a minor class of EGFP-positive cells within the notochord, myotome, and melanocytes, typical of misexpression that occasionally occurs in F0 transgenics (Mosimann et al., 2013). However, a subset of invaded cells that we scored had a fragmented morphology, characteristic of apoptotic cells (Fig. 2C), and were caspase-3-positive (data not shown). Moreover, by 5 dpf, very few EGFP-KRas^V12^ cells remain in these fish, suggesting that these internalized cells eventually die (Fig. 2 D &E). Because cancers do not typically result from a single oncogenic mutation, we sought to test the fate of EGFP-KRas^V12^ cells that also contain a common collaborating mutation in p53. p53 is a pro-apoptotic gene that is frequently mutated in KRas-driven pancreas, colon and lung tumors (Hingorani et al., 2005; Moore et al., 2003; Scarpa et al., 1993), we hypothesized that p53 loss in our model would attenuate death of invading cells.

**Figure 2.**
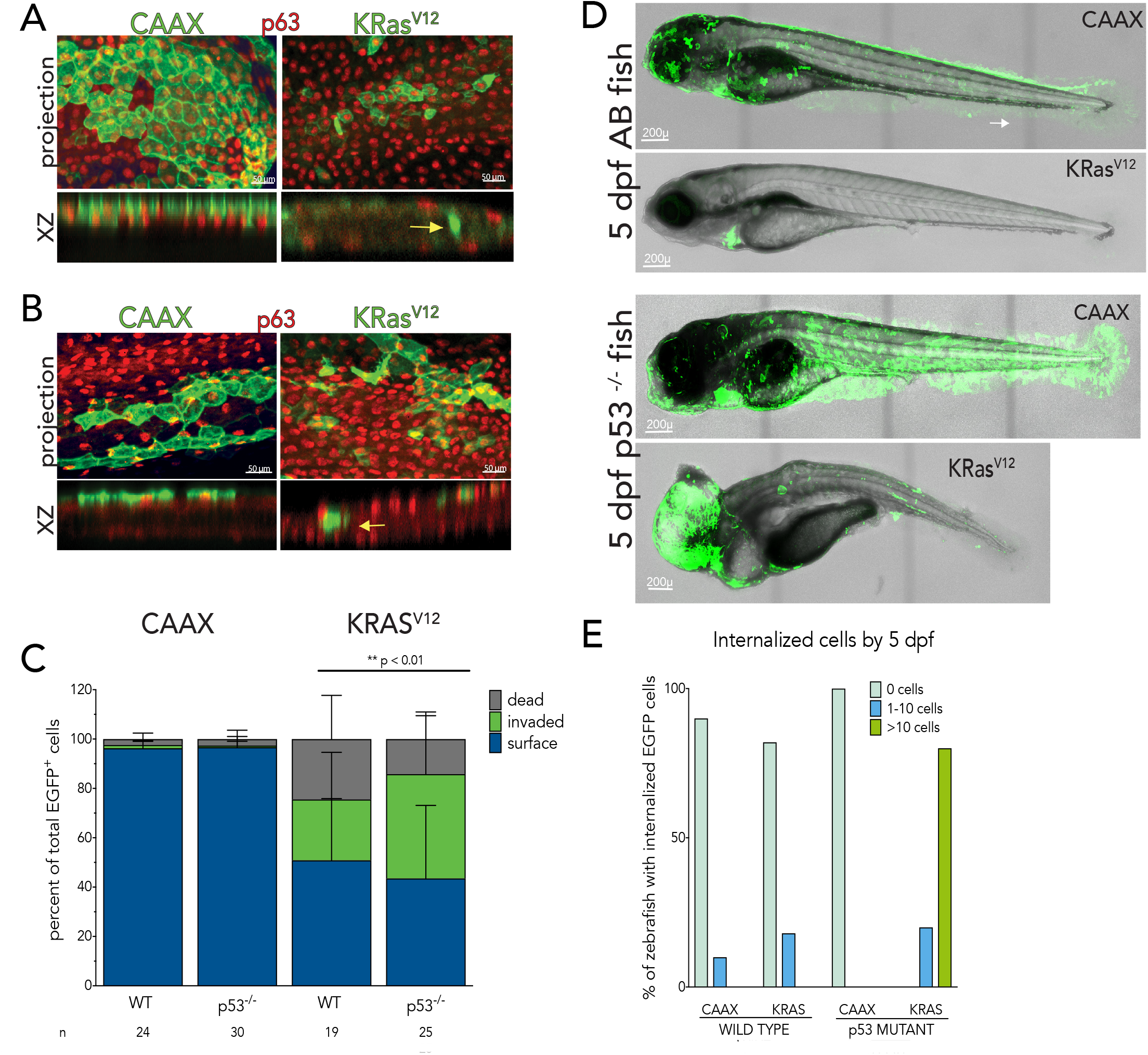
p53 loss increases survival of invading KRas^V12^ cells in the zebrafish body. (A, B) An orthogonal slice view below each projection confocal image shows EGFP-KRas^V12^ cells internalized within the fish body, while CAAX control cells remain along the epidermal surface in AB wild type fish (A) and in p53^-/-^ mutant fish (B). (C) Quantification showing that there are significantly more live internalized EGFP-KRas^V12^ cells upon p53 loss at 48 hpf. (D) p53 mutant zebrafish expressing EGFP-KRas^V12^ contain many internalized cells and masses at 5 dpf whereas wild type zebrafish injected with the same construct lose all cells containing this construct (except the green heart, which indicates trangenesis from Tol2 kit). GFP-CAAX-injected into either wild-type or p53 mutants remain covering much of the epidermis without any cells internalized. (E) Quantification of the percentages of 5 dpf fish containing different numbers of internalized GFP cells.

### Loss of p53 enhances survival of invading KRas^V12^ cells

We found that more KRas^V12^ cells invade and survive when expressed in tp53^zdf1/zdf1^ mutant (hereafter denoted p53 mutant) embryos than wildtype embryos, confirming our hypothesis that p53 loss promotes survival after cells extrude (Fig. 2 B&C). Additionally, more KRas^V12^ cells survive in p53 mutant larvae after 5 dpf, often forming internalized masses in the head (Fig. 2 D&E). This suggests that while KRas^V12^ cells can invade by basal extrusion, they normally die after they invade unless they also contain a collaborating mutation in p53. Thus, to investigate the potential of KRas^V12^ to enable sustained survival of invaded cells, we next analyzed the behavior of KRas-transformed cells lacking p53.

To test the ability of KRas^V12^ to basally extrude and form masses in zebrafish that lacked p53, we co-injected either EGFP-CAAX or EGFP-KRas^V12^ with a p53 translation-blocking morpholino (Robu et al., 2007) so that we could examine extrusions using actin (krt4:Lifeact:mCherry) and other reporter lines. Injection of EGFP-KRas^V12^ with p53 morpholino resulted in significantly more cell masses and extruded at much higher rates than EG-FP-CAAX (SFig. 1 A-C). Unexpectedly, p53 loss randomized the direction of extrusion, resulting in nearly equal numbers of apical and basal extrusions (S. Fig. 1C). The reduction of basal extrusions may be due p53 loss mitigating the autophagy induced by KRas^V12^ mutation (Scherz-Shouval et al., 2010), which disrupts apical extrusion by degrading S1P (Slattum et al., 2014). Another hypothesis is that apical extrusion may act as a fail-safe mechanism to eliminate genetically defective cells when p53 is absent, as we find that CAAX cells have higher rates of extrusion in p53 mutant than in wild type zebrafish (compare Fig. 1E S.Fig. 1C). Although p53 loss decreased the rate of KRas^V12^ basal extrusion, it increased survival of cells that internalized (Fig. 2C) and their ability to survive as internalized masses at 5 dpf (Fig. 2E).

### KRAS cells lose to wild-type cells

Interestingly, although EGFP-KRas^V12^ cells form masses, they lost to non-expressing epidermal cells when compared to EGFP-CAAX control cells in a p53 mutant background (Fig. 2 A and SFig. 1 A), similar to that seen in a wildtype background (Fig. 1A). Quantification shows that 48 hpf zebrafish had far more EGFP-CAAX cells (~100 to 1000) than EGFP-KRas^V12^ cells (~1 to 100) (Fig. 3A). Yet, of the many EGFP-CAAX cells, very few were internalized compared to EGFP-KRas^V12^ cells. Because KRas^V12^ cells extrude at much higher rates than CAAX and other non-expressing cells at this stage in development (Figure SFig. 1C), we surmised that extrusion, at least in part, was responsible for the comparative loss of KRas^V12^ cells to wild type cells in the epidermis.

**Figure 3.**
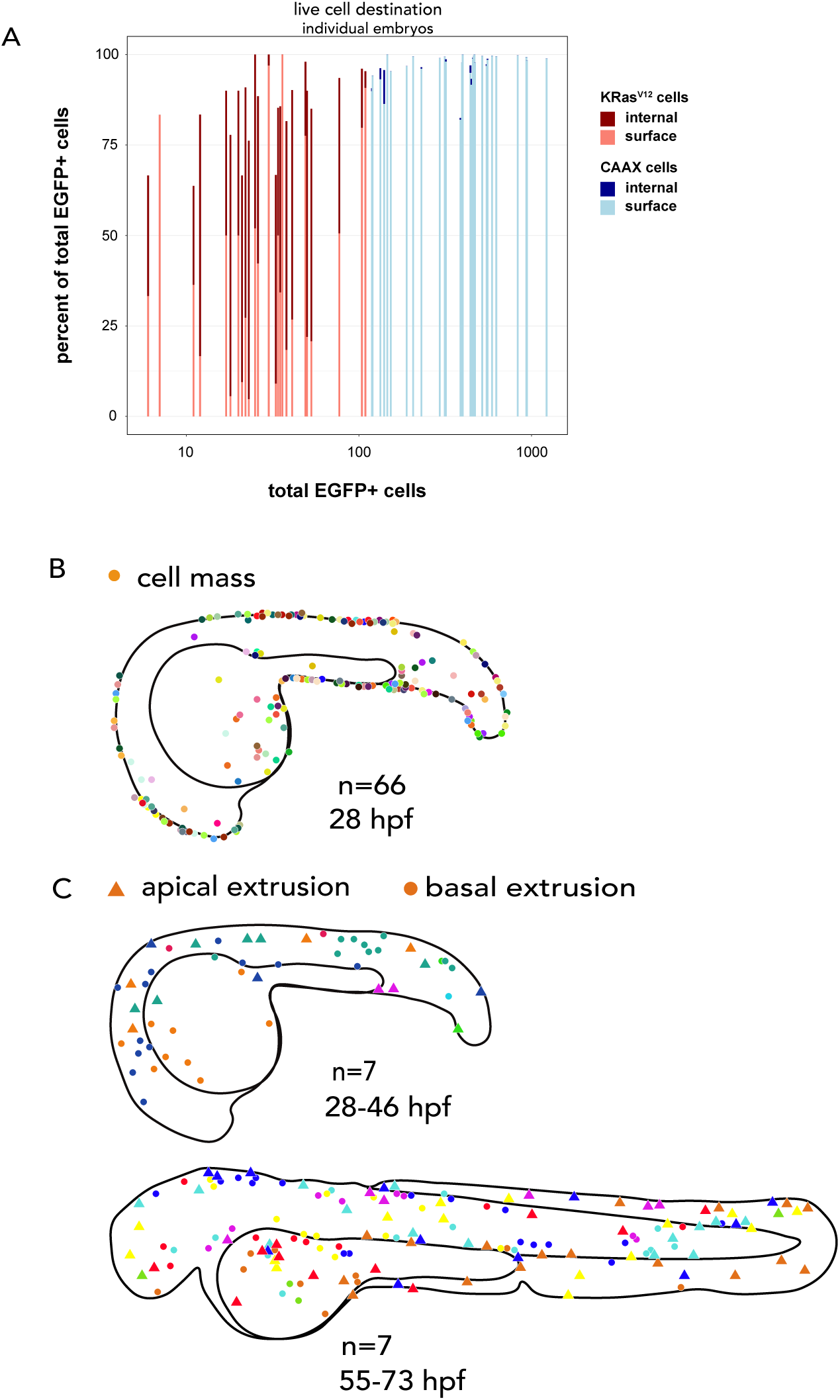
KRas^V12^ cells form masses and basally extrude at distinct sites. (A) Percentages of live cell destination of individual are plotted as a function of the total EGFP+ cells per animal on the X-axis (log scale). There are more internalized KRas^V12^ cells than CAAX cells despite many more epidermal CAAX cells. Maps of where cell masses occur (B) and where apical and basal extrusions occur (C) indicate that these events happen at completely different sites. In both maps, different colors represent different fish analysed.

### KRAS cells form masses and basally extrude at separate sites

Importantly, KRas^V12^ cells extrude at completely separate sites to where they form masses. Masses generally formed at the fin edges (Fig. 3B), sites where cells typically converge, crowd, and extrude in wild-type larvae (Eisenhoffer et al., 2012). The masses at edges were reminiscent of where masses formed in S1P_2_ mutant zebrafish larvae (Gu et al., 2015) and in Piezo1 morphants (Eisenhoffer et al., 2012) when apical cell extrusion is blocked, suggesting a common mechanism for mass formation. However, apical and basal extrusions occurred in central epidermal regions (Fig. 3C), sites where extrusions rarely occur in wild-type fish of this stage. While we quantified extrusions from movies during two time-frames, 28-46 and 55-73 hpf, we only quantified masses prior to filming at 28 hpf, as many of the masses resolve by 72 hpf. The resolution of masses is transient, however, as more develop by 5 dpf (see Fig. 2E). These results suggested that KRas^V12^ expression causes (1) cells to accumulate into masses at sites where they normally extrude and (2) cells to lose to wild-type neighboring epidermal cells by both apical and basal extrusion, the latter of which leads to higher frequencies of invading cells.

### Invading KRas cells deform nuclei and express EMT markers

We next identified and characterized the behaviors of internalized KRas^V12^ cells. We first tested if internalized EGFP-KRas^V12^ cells retained their epithelial identity by staining for E-cadherin. Surprisingly, we found that all internalized EGFP-KRas^V12^ cells lacked E-cadherin (340 cells from 25 fish), whereas surface cells in the same embryos were strongly E-cadherin-positive (Fig. 4A). Additionally, the very few internalized CAAX cells (43 cells from n = 30 fish) were all also E-cadherin-negative (Fig. 4B). However, it was not clear whether loss of E-cadherin was sufficient to switch on mesenchymal markers, predicted during EMT. Therefore, we immunostained EGFP-CAAX and EGFP-KRas^V12^-injected p53 morphants for the mesenchymal marker, N-cadherin. Surprisingly, only some internalized EGFP-KRas^V12^ cells expressed N-cadherin by 48 hpf (Fig. 4 C & D).

**Figure 4.**
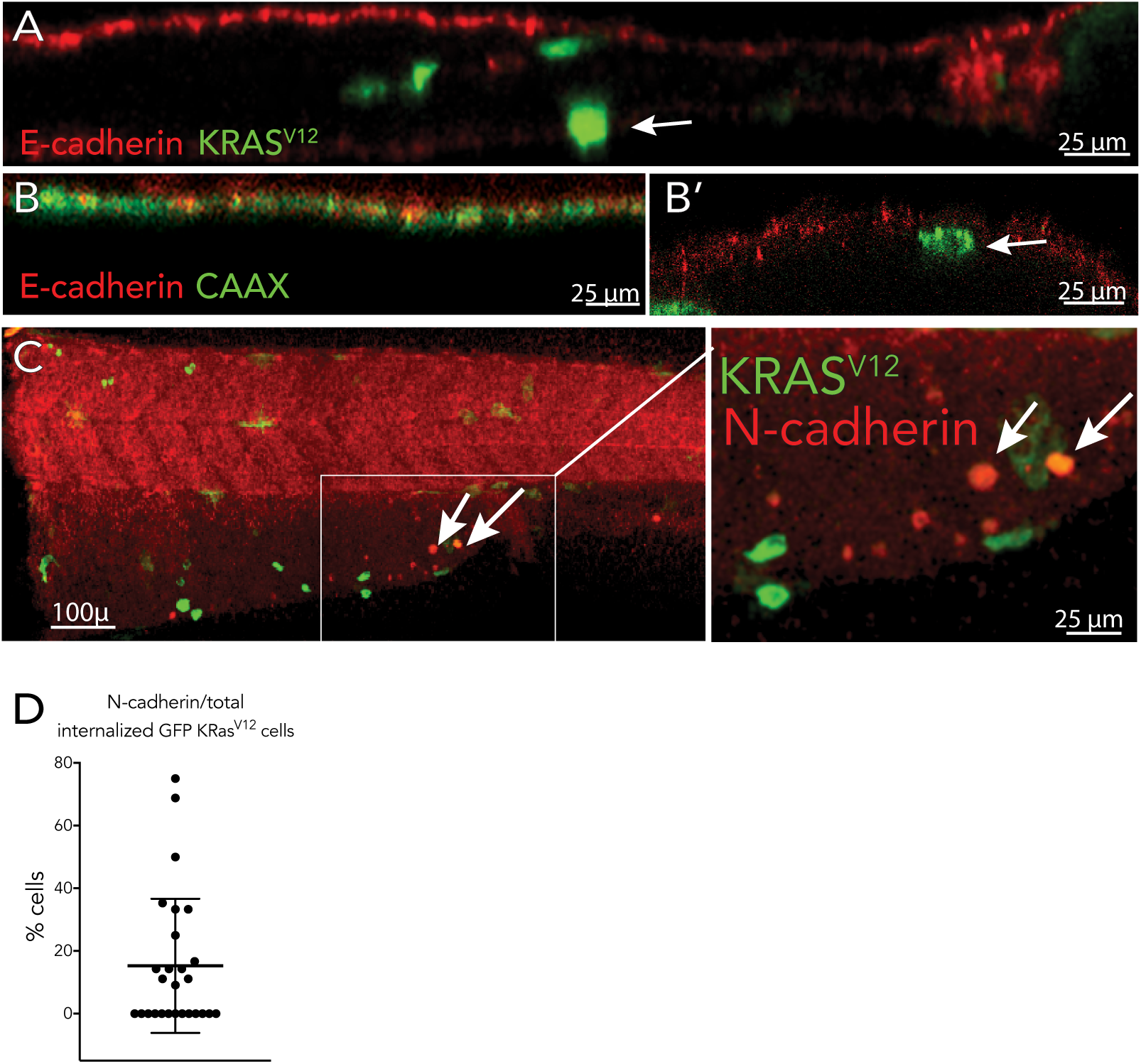
Internalized KRasV12 cells lack E-cadherin but only some express N-cadherin (A) GFP-KRasV12 cells that are internalized lack E-cadherin whereas those at the surface are all E-cadherin-positive. Few GFP-CAAX cells internalize (B) but those that do are also E-cad-herin negative (B’). Only a fraction of internalized cells expresses the mesenchymal marker N-cadherin (C, C’), quantified in (D).

The small percentage of E-cadherin-negative/N-cadherin-positive internalized cells suggested that either expression of mesenchymal markers requires time following invasion or that N-cadherin expression may require other factors. One possible factor that may control epithelial cell de-differentiation could arise from nuclear reprograming by their mechanical contortion as cells migrate through confinement. We noticed that internalized migrating EGFP-KRas^V12^-expressing cells in an H2ava:H2ava-mCherry zebrafish line showed very dim Histone H2A expression, compared to cells within the epidermis (Fig. 5A, SMovie 5). Instead, the nuclei of these cells were diffusely red and deformed, suggesting that as internalized KRas^V12^ cells squeeze through the zebrafish body, their nuclei could become compressed or contorted. Indeed, a movie (Fig. 5B, SMovie 6) of a cell extruding and then migrating in a H2ava:H2ava-mCherry zebrafish line shows that over 42 hours, its H2A-positive nucleus becomes more diffuse and dim as the cell migrates away. Other studies now suggest that differential stiffness and pressures on nuclei can directly affect the fate and identity of the cell (Ivanovska et al., 2015; McGregor et al., 2016; Swift et al., 2013), suggesting that the environment may impact the fate of the invaded cell.

**Figure 5.**
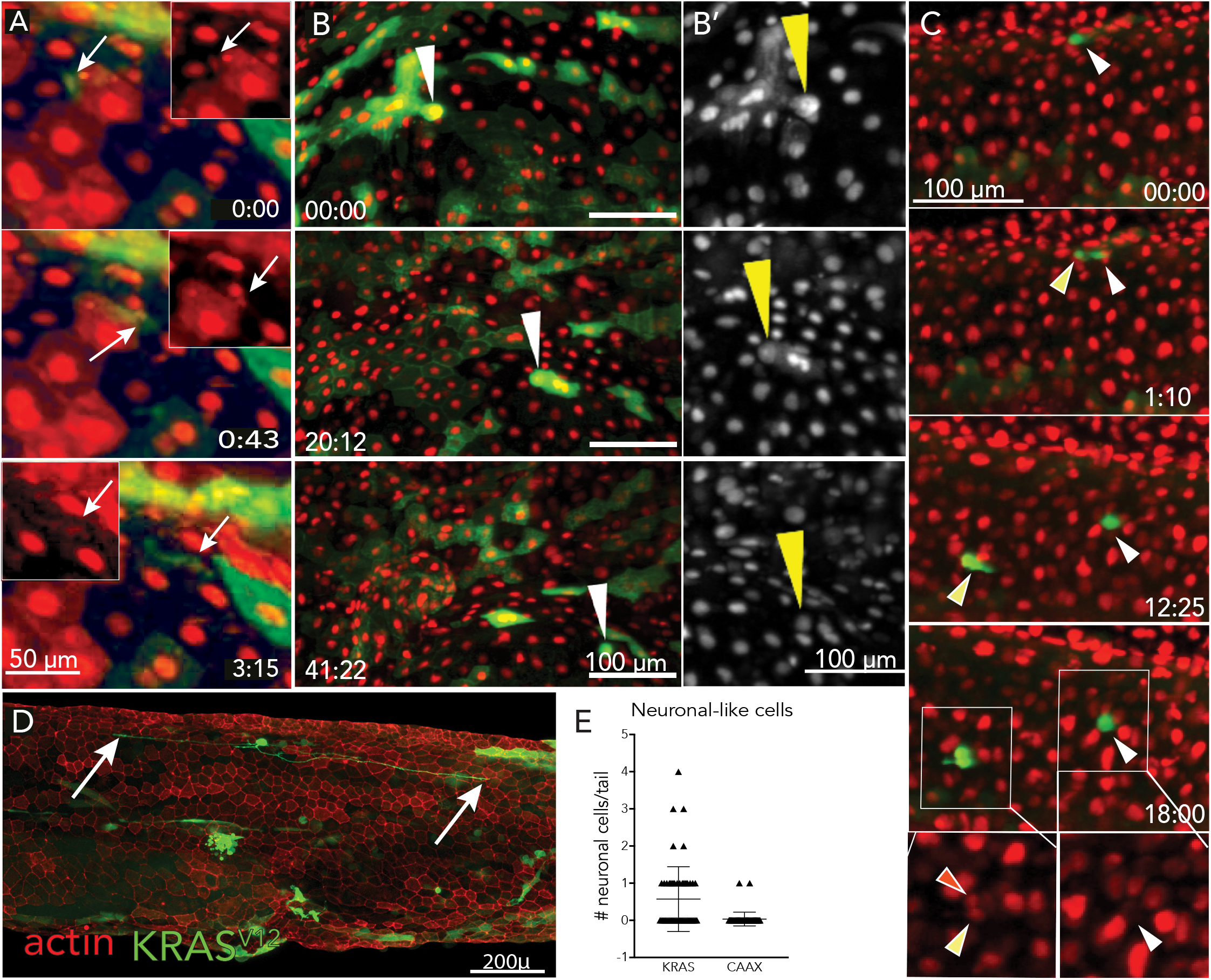
Invading KRas^V12^ cells can divide, contort DNA, and become neuron-like. (A) Stills from a movie (SMovie 5) of migrating internalized EGFP-KRas^V12^ cells (green) that express Histone H2A-mCherry (red) showing that migrating cells have reduced H2A expression and contorted nuclei (arrows). (B) Stills from SMovie 6 show that over time, a cell that extruded expressing Histone H2A-mCherry gradually dims over time as it migrates throughout the zebrafish, with zoomed insets of nuclei only in B’. (C) Internalized EGFP-KRas^V12^ cell (white arrows) from SMovie 7 divides twice and daughter cells migrate away. For A-C, EGFP-KRas^V12^ cells are green and Histone H2A-mCherry are red. (D) Some internalized cells migrate out bidirectionally (also see SMovie 8), similar to neurons (arrows indicate each end of the cell). For all movie stills, time is (hh:mm). (E) Graph showing the number of neuronal-like EGFP-KRas^V12^ versus EGFP-CAAX cells per fish tail, where P-value from unpaired T-tests compared to control is ^****^<0.0001.

Invading KRas^V12^ cells also acquired different fates and characteristics, very rarely seen in control CAAX-injected fish. Frequently internalized, migrating EGFP-KRas^V12^-expressing cells divide and then continue moving in different directions. Fig 5C (SMovie 7) shows a cell dividing twice within a span of 12-15 hours, similar to the division rate of some epidermal cells during this stage of development. Surprisingly, we frequently saw neuronal-like EGFP-KRas^V12^ internalized cells in fixed embryos (32 within 64 confocal projections) and in movies elongating bi-directionally (Fig. 5D & SMovie 8). By contrast, EGFP-CAAX-expressing fish had only two neuronal-like cells from 57 confocal projections of zebrafish tails, suggesting that these were unlikely to arise merely from mis-expression of the construct. Thus, cells that invade by basal extrusion can migrate throughout the body of the zebrafish, divide, and become mesenchymal or other cell types over time.

### EGFP-KRas^V12^ cells circulate in the bloodstream

We next investigated whether basally extruded KRas^V12^-expressing cells enter the vasculature. Using a kdrl:mCherry zebrafish line to label the vasculature, we found that of 49 p53 morphants filmed, ~72% had GFP-KRas^V12^ cells circulating within the bloodstream (Fig. 6A, SMovie 9). By contrast, only one of 175 GFP-CAAX p53 morphants had circulating GFP-cells. Interestingly, we found that 65% of 42 wild-type AB zebrafish also had circulating EGFP-KRas^V12^ cells (Fig 6B). Presumably, many of the KRas^V12^-expressing cells die in a wild-type background, as we did not see any EGFP-cells at 5 dpf. These results suggest that basally extruded KRas^V12^-expressing cells can intravasate into the vasculature even at early stages yet only persist in a p53 mutant. Additionally, in movies using the kdrl:mCherry line and another line that expresses mCherry in red blood cells, Gata1a:mCherry, we found that circulating GFP-KRas^V12^ cells frequently impeded the blood flow for long intervals (Fig. 6C & D, SMovie 10 & 11).

**Figure 6.**
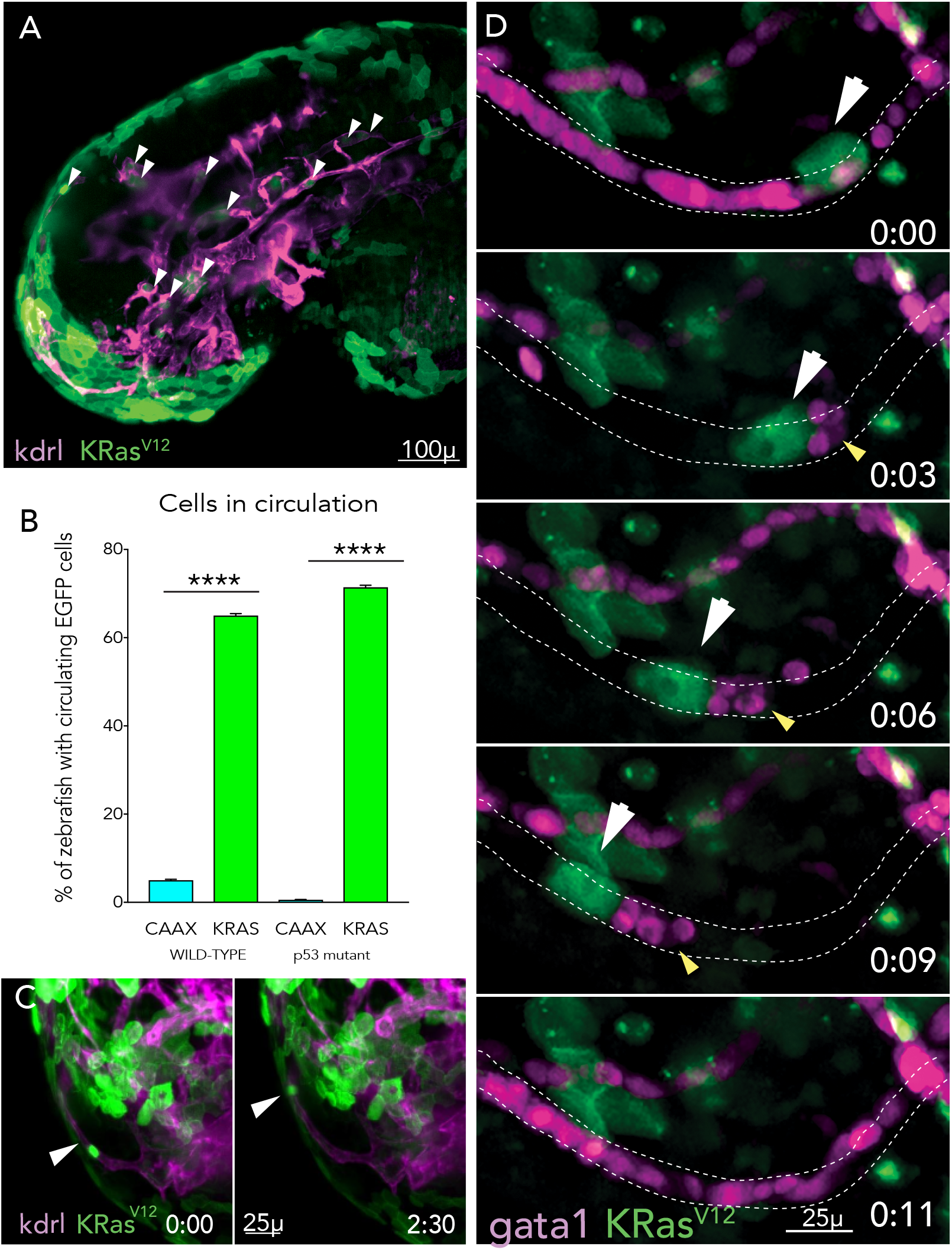
Invading KRas^V12^ cells enter the bloodstream and can impede blood flow. (A) Cross section from SMovie 9 showing numerous EG-FP-KRas^V12^ cells within blood vessels. (B) Quantification of zebrafish containing circulating EGFP-KRas^V12^ versus EG-FP-CAAX cells in wild-type or p53 mutant backgrounds. P-values from unpaired T-tests compared to control are ^****^<0.0001, where error bars = s.e.m. (C) Stills from SMovie 10 showing GFP-KRas^V12^ cells in a small vessel. (D) Stills from a time-lapse movie (SMovie 11) of an EGFP-KRas^V12^ cell that is trapped within a blood vessel, blocking blood cell flow and eventually being carried out presumably by pressure from the blood flow. For all movie stills, time is (hh:mm).

### Basally extruding cells simultaneously lose epithelial identity via membrane shedding

Despite the fact that we routinely saw high rates of basal extrusion and cell internalization in EGFP-KRas^V12^-expressing zebrafish, frustratingly, we could never capture these two events consecutively in our movies. This was because KRas^V12^ cells typically lost their GFP signal as they basally extruded. To better resolve how this GFP signal was lost, we expressed EGFP-KRas^V12^ in ionocytes (using ionocyte:Gal4 x UAS:NTR-mCherry lines, injected with UAS:EGFP-KRas^V12^) that are sparsely distributed throughout the epidermis, so that neighboring cells did not obscure resolution of the extruding cells. We observed a startling discovery: Ionocytes expressing EGFP-KRas^V12^ (tethered to apical membrane) and NTR-mCherry (in the cytoplasm) split into a red and a green fragment, as it basally extrudes (Fig. 7A and SMovie 12). The larger red cell fragment then migrates away from the site of extrusion. The loss of apical membranous GFP during basal extrusion was not a feature limited to ionocytes. Outer epidermal cells expressing EGFP-KRas^V12^ and cytoplasmic mCherry (periderm:Gal4; UAS:N-TR-mCherry injected with krt4:EGFP-KRas^V12^) also lose their apical GFP as they basally extrude so that a red cell migrates away (Fig. 7B and SMovie 13). From these results, we hypothesized that the apical membrane, with the associated EGFP-KRas^V12^, of extruding cells may be stripped off mechanically during basal extrusion.

**Figure 7.**
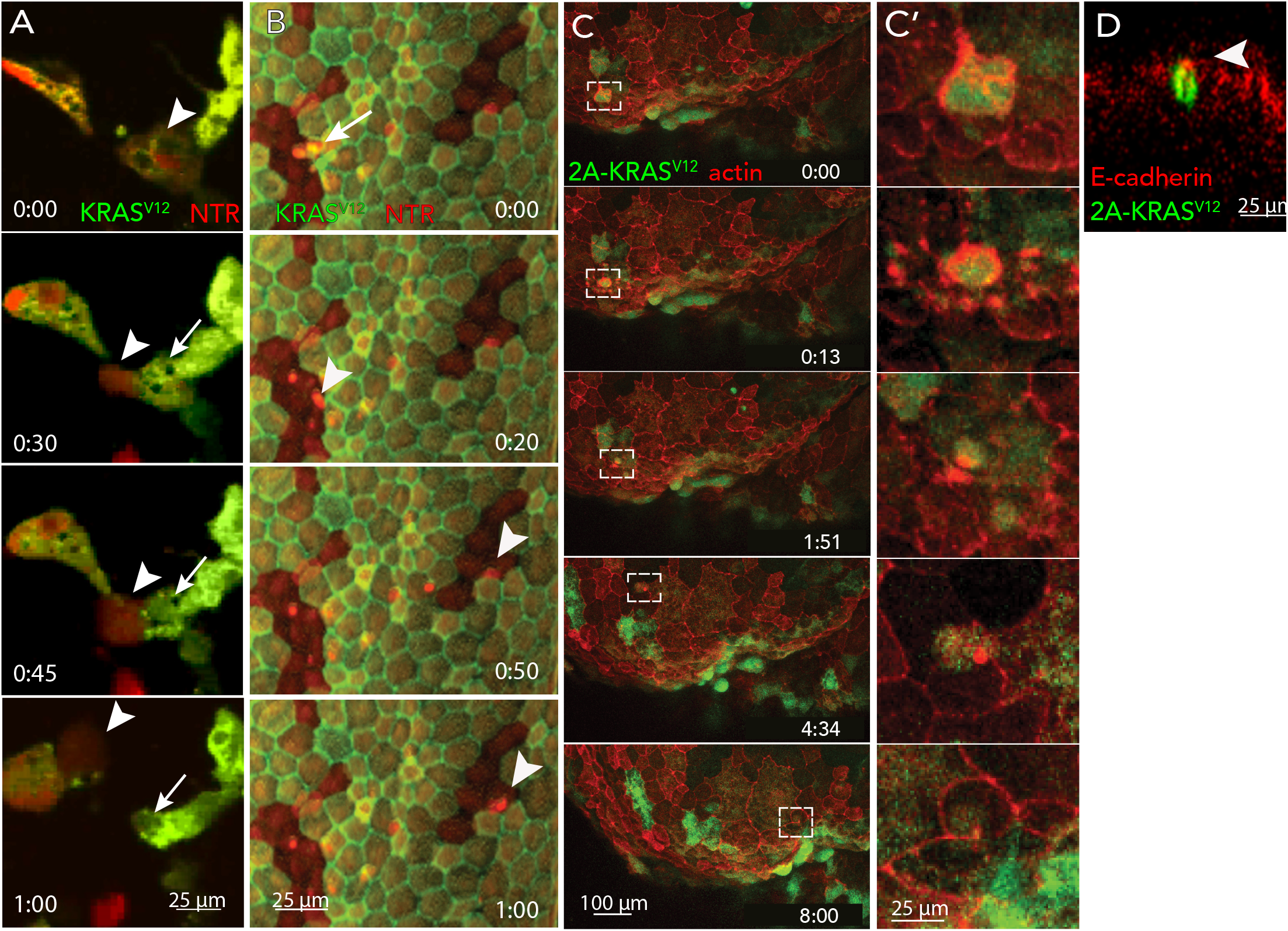
Basally extruding cells scrape off apical membranes and E-cadherin. (A) Stills from SMovie 12 where an EGFP-KRas^V12^ ionocyte cell expressing NTR-mCherry in the cytoplasm shows that as the cell basally extrudes, it pinches off the EGFP within the apical membrane (arrow). Then the remaining red cell (arrowhead) migrates away. (B) Stills from SMovie 13 where an EGFP-KRas^V12^ cell also expressing GFP-Lifeact and NTR-mCherry shed EGFP (arrow) and migrate away as a red cell (arrowhead). (C) Stills from SMovie 14 where GFP is expressed separately to KRas^V12^ so that it labels the cytoplasm shows a portion of the cell extrudes apically while the remainder migrates away with actin (red) at the back of the cell. (C’) insets showing magnifications of the cell. For all movie stills, time is (hh:mm). (D) XZ picture of a KRas^V12^-cell where EGFP is in the cytoplasm show that E-cadherin is constricted apically into one point as it basally extrudes. Scale bars = 10μm.

To test this possibility, we cloned a T2A self-cleaving peptide between EGFP and KRas^V12^, allowing EGFP to mark the cytoplasm of invading cell, rather than its apical membrane and expressed this construct in a line that labels epidermal actin in red (krt4:Lifeact:mCherry). This new construct allowed us to visualize EGFP in a single cell before, during and after the extrusion. EGFP-T2A-KRas^V12^ cells revealed that as cells basally extrude, they shed their apical membranes and bleb extensively, similar to a cell undergoing apoptosis (Fig. 7C and SMovie 14). Following extrusion, the remainder of the cell migrates throughout the body of zebrafish over 8 hours (SMovie 14). Interestingly, here, and in other movies, the apical actin “scar” from the basal extrusion remains at the back of the migrating cell, suggesting that it uses a ‘stable-bleb’ type of rapid motility, characteristic of cells under low adhesion and high confinement (Liu et al., 2015a; Ruprecht et al., 2015a; Welch, 2015b). We found that all KRas^V12^ cells shed their apices as they basally extrude and invade in over 50 other movies. Importantly, immunostaining an EGFP-KRas^V12^ cell in the middle of extrusion shows E-cadherin becomes constricted into an apical point with the rest of the cell beneath it (Fig. 7D). Together, our findings suggest that basal extrusion drives invasion of KRas^V12^ cells and simultaneously strips off their apical surfaces, and with them, their E-cadherins, essential to epithelial function. While EGFP-KRas^V12^ expression returns later, E-cadherin does not.

## DISCUSSION

Here, we present a new model for tumor cell invasion: oncogenic KRas signaling hijacks a mechanism normally used to promote cell death to instead drive cells to invade and simultaneously lose their epithelial identity (graphical abstract). Wild-type epithelial cells extrude out apically to die when they become too crowded. However, KRas^V12^-transformation disrupts apical extrusion, causing cells to instead form masses at sites of crowding or aberrantly extrude basally at independent sites. Here, we find that apical constriction not only extrudes the cell basally but also scrapes off the entire membrane apical to the constricting junctional actin and, with it, E-cadherin. Once cells invade by extrusion, they migrate throughout the rest of the body, divide, become different cell types, and intravasate into the bloodstream. Because apical extrusion is disrupted in many invasive cancer types, we predict that this new invasion mechanism revealed in our zebrafish model will drive initiation of metastasis in cancers with similar signaling. Our findings also suggest that although KRas mutations are generally considered early mutations in pre-cancerous lesions, they impart an intrinsic metastatic propensity. However, disseminated tumor cells only survive when anoikis is blocked by collaborating mutations in p53 or other pro-apoptotic factors.

### New mechanical mechanism for epithelial to mesenchymal transition

Using zebrafish embryonic epidermis to model simple epithelia, we can for the first time visualize transformed cells invading live from the epithelia where carcinomas originate. This has enabled us to discover a new rapid, mechanical mechanism for EMT that occurs at sites independent of a primary mass. This mechanism lies in stark contrast to the prevailing EMT model, where cells invade from primary tumors after transcriptionally downregulating E-cadherin via developmental signaling pathways (Hanahan and Weinberg, 2011; Kalluri and Weinberg, 2009). While the established EMT model can also initiate tumor spread, we think that its occurrence may be overestimated for the following reasons: (1) pathology slides are made from primary tumor sites, rather than distant sites where cells may be basally extruding, and (2) most metastasis studies in mice follow cells migrating from a transplanted, pre-formed mass. Interestingly, from all our movies (~600), we witnessed only one case where EGFP-KRas^V12^-positive cells escape and migrate from a mass (Smovie 15).

Our live imaging revealed that cells invade by basal extrusion at sites completely unrelated to primary masses. Moreover, as cells invade by basal extrusion, they rapidly and mechanically scrape off their EGFP-KRas^V12^, E-cadherins, and other apical membrane proteins and lipids. The EGFP-KRas^V12^ is only transiently lost, as we later see EGFP-positive cells within the body of the fish. However, E-cadherin disappears permanently in single invading cells. Loss of E-cadherin may be due to protein degradation when not engaged with other extracellular E-cadherins, as seen previously (Adams et al., 1998). Regulation of E-cadherin expression by protein stability, rather than transcriptional downregulation, could account for how quasi-mesenchymal tumor cells can transition back to epithelial, should they encounter more epithelial cells. This mechanical EMT could account for how tumor cells typically retain a mix of epithelial and mesenchymal markers, unlike cells during normal development (Ye and Weinberg, 2015).

Importantly, the oncogenic mutations in KRas and p53 used in our studies are critical drivers for aggressive, invasive tumors with poor prognosis, including pancreatic, kidney, and many lung and colon cancers. Although the transient overexpression of KRas^V12^ in our zebrafish assay may not perfectly model the activity of KRas in these aggressive cancers, the ability of these epidermal cells to form masses and basally extrude mimics the activity of pancreatic tumor cells both in vitro and in vivo (Gu et al., 2015; Hendley et al., 2016). Furthermore, classic transcriptional drivers of developmental EMT are dispensable for metastasis in pancreatic cancer (Zheng et al., 2015) and lung cancer (Fischer et al., 2015). Therefore, we predict that the cellular behavior that drives this aggressive tumor type in the fish will be conserved in human cancers.

### Implications for mechanical EMT

Our findings that basal extrusion and sudden loss of the entire apical membrane can drive tumor invasion leads to numerous implications for metastatic tumor cells. Many of these changes may address perplexing clinical findings and could shape the way we target metastatic tumors.

#### Blebbing cell migration following extrusion

As cells basally extrude and invade, the remnants of the apical constricted actin ring remained at the rear of cell as they migrate, which could propel its migration throughout the body. This type of motility is consistent with the recently discovered, rapid, contractile ‘stable bleb’ cell migration that predominates in low adhesion and high confinement environments (Liu et al., 2015b; Ruprecht et al., 2015b; Welch, 2015a). Initially, the migration behavior of internalized KRas^V12^ cells raised concerns that they had either become macrophages or that macrophages had ingested them, however, immunostaining revealed no co-localization (S. Fig. 2). Thus, their similar motility is likely due to the findings that both tumor and immune cells use a stable bleb migration (Welch, 2015a). In our experiments, the apical actomyosin constriction of the basally extruding cell may initiate stable bleb migration by polarizing actin and myosin at the cell rear. Stable bleb motility would be reinforced as cells invade through confined cellular and extracellular structures with reduced substrate contact.

#### Loss of apical membrane

The sudden loss of E-cadherin and other apical membrane components could greatly impact the identity of invaded cells and our ability to diagnose and target them clinically. For instance, if most metastatic tumor cells lose apical cell surface markers, methods to identify circulating tumor cells that rely on membrane-associated markers like EpCAM may be ineffective (Gabriel et al., 2016). Additionally, mechanical apical membrane shedding could also eliminate receptors such as EGFR or ERBB2, rendering cancer therapies targeted to these receptors ineffective (Meyer et al., 2013; Wilson et al., 2018). Loss of E-cadherin is a common mechanism by which cells can escape anoikis, which is critical for metastatic tumors to survive at distant sites (Fouquet et al., 2004; Frisch et al., 2013; Kumar et al., 2011). Finally, this virtual cytokinesis of the cell apex could instantly depolarize the invading cell and loss of cell polarity is a known hallmark of cancer initiation and progression (Gabriel et al., 2016; Muthuswamy and Xue, 2012; Saito et al., 2018). Further studies will need to follow the fate of these invading cells and what different programs they activate and de-activate.

#### Mechanical impact on nucleus and gene expression

Our movies of zebrafish expressing Histone H2A-mCherry by its endogenous promoter suggest that nuclei of migrating KRas^V12^ cells become highly contorted during migration (Fig. 5A&B). Mechanical stiffness and tensions can impact not just nuclear shape changes but also regulate transcription of genes controlling cell differentiation and proliferation (Belaadi et al., 2016; Elosegui-Artola et al., 2017; Swift et al., 2013). Mechanical stretch can directly allow YAP into the nucleus by expanding pore entry, thereby enabling cell proliferation (Elosegui-Artola et al., 2017). Alternatively, the cell environmental stiffness can impact cell fate (Swift et al., 2013). Thus, while previous studies have assumed that EMT programs were initiated by transcription factors that drive cell invasion by first down-regulating E-cadherins, our new mechanical model of EMT suggests that cells first invade by basal extrusion and change their transcriptional programs and cell fates later. Because only some of the invaded cells had mesenchymal markers, one possibility is that mechanical contortion of the nucleus during migration or differential environmental stiffness could dedifferentiate epithelial cells after they invade. Another factor may be due to the environment the invaded transformed cell contacts.

#### Transdifferentiation of KRas^V12^ cells

Internalized KRas^V12^ cells had numerous different fates in our movies and fixed embryos. While some cells could continue to divide, others appeared to become different cell types. Based on the fact that our experiments use zebrafish embryos, the different cell morphologies could reflect the fact that these cells are still plastic during development. On the other hand, pancreatic cancer, which is driven by the same mutations in our study, is notably highly stromal with neuronal infiltration (Rasheed et al., 2012). Additionally, many aggressive cancers recruit neurons and rely on them for their growth (Demir et al., 2015; Saloman et al., 2016). Cre-labeling pancreatic epithelial cells from a genetically engineered mouse model for pancreatic cancer showed that the stromal component derives from epithelia, rather than being recruited, as previously expected (Rhim et al., 2012). Our finding that invading KRas^V12^ cells frequently become neuron-like suggests that these cells may re-differentiate into neurons, rather than recruit them.

Invasion occurs at sites distinct from cell masses It has long been assumed that metastases derive from a tumor mass; however, our data demonstrate that tumor cells invade at sites completely independent to a primary mass. Normally, zebrafish epidermal cells proliferate near the midline and then migrate out to the fin edges where they converge and extrude (Eisenhoffer et al., 2012). When apical extrusion is blocked by expression of KRas^V12^, cells form masses at the fin edges where wild-type cells would have extruded. Additionally, KRas^V12^ cells extrude predominantly basally in regions where wildtype epidermal cells normally proliferate. This suggests that KRas^V12^ cells lose to wild-type cells by epithelial extrusion but because many extrude basally and survive, they ultimately ‘win’ at other sites within the zebrafish. Our finding that cells invade by basal extrusion at times and sites independent to where masses form suggests that tumor cell dissemination can occur independently of tumor formation, similar to the findings by (Rhim et al., 2012). In fact, ‘cancer of unknown primary site’ syndrome is not uncommon (Neben et al., 2008) and may result through this mechanism.

#### Early circulating tumor cells

Our findings that EGFP-KRas^V12^ cells enter the bloodstream rapidly, without p53 mutations, suggest that cancers driven by KRas could be particularly invasive at the outset. One caveat of our finding is that it may be much easier to enter the bloodstream when the vasculature is still developing in an early embryo than in an adult human. However, a genetically engineered pancreatic cancer mouse model found circulating tumor cells before detectable tumors (Rhim et al., 2012), similar to our findings. Further, the fact that pancreatic cancer is rarely diagnosed in humans without metastases also suggest that mutations in KRas and p53 that drive these tumors are intrinsically invasive (Das and Batra, 2015). The fact that EGFP-KRas^V12^ cells can intravasate into vessels and block blood flow in smaller vessels may explain why blood clots are associated with cancer metastasis and commonly occur before a cancer diagnosis, especially in pancreatic cancer (Boone et al., 2018; Kang and Tang, 2012). Because a KRas mutation alone is not sufficient to cause cancer, but can enable cells to enter the circulation, blood tests could potentially screen for people who are at high risk for these cancers. If so, chloroquine, which targets KRas-mutant cells with high autophagy (Boone et al., 2018; Kang and Tang, 2012), may provide a putative, preventative therapy.

#### New model to screen for targets that kill metastatic tumors

Our zebrafish assay to investigate how tumor cells invade should also provide an excellent model to screen for new drugs or drug combinations that specifically target disseminated tumor cells. Metastasis is the predominant reason patients succumb to their cancers, emphasizing our current inability to target metastatic tumors. Through the ability to follow all steps of metastasis in our zebrafish model, we should be able to learn how cells disseminate and why they overtake certain organs. Moreover, because we can readily see every single disseminated cell, we should be able to screen for drugs that kill every tumor cell in our zebrafish assay. This may better inform us why current treatments have been ineffective and pave the way for treating metastatic disease.

## CONTACT FOR REAGENT AND RESOURCE SHARING

Further information and requests for resources and reagents should be directed to and will be fulfilled by the Lead Contact, Jody Rosenblatt.

## EXPERIMENTAL MODEL AND SUBJECT DETAILS

The AB wildtype line was obtained from the University of Utah Centralized Zebrafish Animal Resource (CZAR). The tp53^zdf1/zdf1^ was a kind gift from Rodney Stewart. The enhancer trap lines were generated in the labs of Chi-Bin Chien and Richard Dorsky (Otsuna et al., Dev Dyn 2015) and previously characterized (Eisenhoffer et al., J Cell Science 2016). All the groups mentioned are at the University of Utah.

Zebrafish lines were maintained at the CZAR in an independent recirculating water system held at 28°C. A 14 h/10 h light/dark cycle was used. The animals were housed in 2.8-L tanks at a density of 20-25 individuals per tank. Two male-female pairs per mating tank were used to generate embryos, which were rapidly collected in E3 embryo water (4.96 mM NaCl, 0.18 mM KCl, 0.33 mM CaCl_2_, 0.86 mM MgCl_2_, pH 7.2) and assigned randomly to experimental groups. All procedures adhered to institutional IACUC guidelines.

## METHOD DETAILS

### Tol2kit cloning for zebrafish lines

All cloning procedures followed protocols detailed in Kwan et al., Dev Dyn 2007, the Invitrogen Gateway Technology Manual, and the Tol2kit wiki (http://tol2kit.genetics.utah.edu/index.php/Main_Page). We cloned the krt4 promoter sequence into p5E (gift from David Grunwald), and human K (in a pEGFP-C3 backbone, KRas^V12^ (isoform 4b, gift from Channing J. Der, University of North Carolina, Chapel Hill, NC) or CAAX (Tol2kit, Kristen Kwan) into pME using the BP Clonase II Enzyme mix (Invitrogen). To generate the krt4:EGFP-KRas^V12^ or krt4:EGFP-CAAX constructs, the following were recombined into pDestTol2CG2 using the LR Clonase II Plus Enzyme mix (Invitrogen): p5E-krt4, pME-EGFP-KRas^V12^ or pME-EGFP-CAAX, and p3E-polyA. The EGFP-T2A-KRas^V12^ construct was synthesized commercially and cloned into pUC57 (GenScript), then sequential BP and LR reactions were done to recombine it into pME and pDestTol2CG2, respectively, as stated above.

Transposase and ⃠-bungarotoxin mRNA were in vitro transcribed using SP6 and T7 mMESSAGE mMACHINE Transcription Kits (Invitrogen), respectively, and purified using NucAway spin columns (Invitrogen). The concentrations of all nucleic acids used were measured using a NanoDrop 1000 (Thermo Fisher) or an EPOCH 2 microplate (BioTek) spectrophotometer.

### Microinjections and sorting

2 nl of a 10-μl injection mix comprised of 100 ng krt4:EGFP-CAAX or 150 ng krt4:EGFP-KRas^V12^ or 150 ng krt4:EGFP-t2a-KRas^V12^, 200 ng transposase mRNA, and 1 μL phenol red (Sigma) in nuclease-free dH_2_O (Ambion) was microinjected (Harvard Apparatus) into each one-cell embryo. Some experiments included 0.2 pmol p53 morpholino (Gene Tools) or 25 ng ⃠-bungarotoxin mRNA. Embryos were sorted for expression of fluorescent transgenes at 1 dpf, dechorionated with forceps at 1 or 2 dpf, incubated in E3 with 0.003% N-phenylthiourea (PTU-E3), and prepared for live imaging or fixed and immunostained.

### Time-lapse confocal microscopy of live embryos

The protocol for mounting embryos for live imaging has been previously described (Eisenhoffer and Rosenblatt, J Vis Exp 2011). Briefly, dechorionated embryos were anesthetized in 0.02% tricaine in PTU-E3 for about 5 minutes or until no visible movement is observed, then suspended in 0.4-1% low-melt agarose. The embryos were mounted in agarose as close as possible to the #1.5 glass coverslip within a slide chamber and then covered with 0.02% tricaine in PTU-E3 and incubated in a controlled environment chamber at 28°C and 85% humidity. We used either an Andor Revolution spinning disk confocal microscope (Nikon CFI Plan Apo 20X/0.75 DIC M ∞/0.17 WD 1.0 objective), a Nikon A1R (Nikon CFI Plan Apo 20X/0.75) resonant scanning system, or a Leica SP8 white light laser confocal.

### SPIM of live embryos

Selective Plane Illumination Microscopy (SPIM) was used for long-term in toto imaging using one of two custom-built SPIM setups: multidirectional SPIM (mSPIM) (Huisken and Stainier, 2007) and four-lens SPIM setup (Schmid et al., 2013). mSPIM was equipped with an UMPlanFL N Olympus 10 x/0.3 NA detection objective, a 488 nm and a 561 nm Coherent Sapphire laser, two Zeiss 10x/0.2 illumination objectives and two Andor iXon 885 EM-CCD cameras. The whole embryo was imaged from several angles with 7x overall magnification and an axial resolution (z-stack spacing) of 3 μm every 5-7 min for up to 3 days. To image the whole embryo, several regions were acquired and stitched together using custom image processing plugins in Fiji (Schindelin et al., 2012), based on a stitching tool (Schindelin et al., 2012). For visualization, maximum intensity projections were generated and registered using SimpleElastix and GUI-based manual rigid registration.

The four-lens SPIM setup consists of four identical water-dipping Olympus UMPLFLN 10x/0.3 objectives, two for illumination and two for detection. Two Toptica iBeam smart lasers were externally triggered for alternating double-sided illumination. The laser beam was split 50/50 and directed onto a continuous running galvanometric mirror (1 kHz, EOPC), which pivots the light sheet and reduces shadowing effects in the excitation paths due to absorption of the specimen (Huisken and Stainier, 2007). Light sheets were generated with cylindrical lenses and projected with telescopes and the illumination objectives onto the focal plane of both detection lenses. The focal planes of the two detection objectives were imaged onto two Andor Zyla sCMOS. The whole embryo was imaged from several angles with an axial resolution of 2 μm every 30 seconds up to 5 minutes for up to 36 hours. A custom LabVIEW (National Instruments) program was implemented to adjust stage positions, stack coordinates and various parameters for time-lapse acquisition. A custom fusion program was used for visualization and generation of maximum intensity projections.

For long-term time-lapse acquisition, embryos were embedded in 0.1% low-melting-point agarose inside fluorinated ethylene propylene (FEP) tubes, essentially as described in (Kaufmann et al., 2012) and (Weber et al., 2014). 0.016% ethyl 3-aminobenzoate methanesulfonate salt was used in the E3-filled-imaging chamber only when bungarotoxin mRNA was not co-injected with DNA constructs.

### Immunostaining and imaging of fixed embryos

The protocol for immunostaining embryos has been previously described (Gu et al., Elife 2015). Briefly, dechorionated and anesthetized embryos were fixed in 4% paraformaldehyde and 4% sucrose + 0.1% Triton X-100 in PBS overnight at 4°C, blocked with 10% goat serum for 1 h, then stained with a cocktail of primary antibodies overnight at 4°C: chicken ⃠-GFP (Abcam, 1:2000), rabbit ⃠-p63 (GeneTex, 1:100), mouse ⃠-E-cadherin (BD Biosciences, 1:200), rabbit ⃠-caspase-3 (BD Biosciences, 1:100), mouse ⃠-N-cadherin (BD Biosciences, 1:100). The primary antibodies were washed off with six 20-minute washes of 0.5% PBST (PBS + 0.5% Triton-X). Embryos were a incubated with a cocktail of goat ⃠-chicken-AlexaFluor488, goat ⃠-rabbit-AlexaFluor^568^, and goat ⃠-mouse-AlexaFluor^647^ (all secondary antibodies at 1:200) in 10% goat serum overnight at 4°C, followed by four 20-minute washes with 0.5% PBST. Nuclei were stained with 1 μM DAPI for 30 minutes and washed twice with 0.5% PBST. The immunostained embryos were taken through a glycerol series (25%, 50%, 70% glycerol in PBS) and finally mounted in 70% glycerol in PBS with a #1.5 glass coverslip. A Nikon A1R galvano scanning system (20X Plan Apo) or a Leica SP8 (20X Plan Apo CS2 0.75), or a Zeiss LSM 880 Airy inverted with a LD LCI Plan-Apochromat 25X 0.8 objective were used to image fixed samples.

## QUANTIFICATION AND STATISTICAL ANALYSIS

1 dpf embryos were anesthetized in 0.02% tricaine and dechorionated to count cell masses manually counted under a brightfield dissection microscope.

Internalized and surface cells were manually classified and counted from confocal images in orthogonal slice projections immunostained for p63 or E-cadherin to define basal or periderm, respectively, of the zebrafish epidermis. Cells were classified as live or dead using caspase-3 immunostaining and general morphology. EGFP fusion proteins misexpressed in muscle, notochord, and pigment cells were excluded from the final analysis. Live cell extrusions were manually quantified from time-lapse movies as apical or basal using orthogonal slice projections.

All statistical calculations were performed using the GraphPad Prism analysis suite or Microsoft Excel functions. We used the unpaired Student t-test assuming Gaussian distribution and unequal standard deviations.

## ACKNOWLEDGEMENTS

We thank Adam Gardner for early studies that were not included in this report; Channing Der, George Eisenhoffer, and David Grunwald for plasmids to construct KRas expression vectors; Kristen Kwan for the CAAX construct, donor vectors, and help with Tol2kit; and Rodney Stewart for the tp53zdf1/zdf1 zebrafish line. We thank Russell Bell for his assistance with graphing internalized cells and Minna Roh-Johnson and James Gagnon for helpful feedback on our manuscript. A National Institute of Health R01GM102169 and a Howard Hughes Faculty Scholar Award 55108560 to J. R., an EMBO long-term fellowship to G. M. S., and P30 CA042014 awarded to Huntsman Cancer Institute core facilities supported this work. We thank the Centralized Zebrafish Animal Resource, Fluorescence Microscopy, and Mutation Generation and Detection Cores in the Health Sciences Cores at the University of Utah and the Light Fluorescent Microscopy Facility at the Max Planck Institute of Molecular Cell Biology and Genetics, Dresden, Germany. An NCRR Shared Equipment Grant #1S10RR024761-01 paid for microscopy equipment.

## Contributions

J. R. designed experiments, interpreted, analyzed data, and co-wrote the manuscript. J. F. made zebrafish Tol2 constructs, designed experiments, interpreted data, performed most live and fixed imaging experiments throughout the paper, and co-wrote the manuscript. G. M. S. developed the first fish expressing KRas and CAAX constructs and performed the initial experiments showing that KRas cells extrude basally in zebrafish and get into the bloodstream, designed experiments, interpreted data, and performed all of the imaging on the light sheet microscope. N. R. and M. F. J. maintained zebrafish lines, injected and immunostained embryos, and quantified many of the experiments. S. D. operated mSPIM and wrote code for image acquisition and visualization. J. H. supported G.M.S and S.D for SPIM imaging. D. H. quantified location of cell masses and extrusions and helped genotype p53 mutants.

## Supplemental Movie Legends

SMovie 1 Krt4-GFP-CAAX cells remain in the epidermis without extruding.

SMovie 2 Krt4-GFP-KRas^V12^ cell apically extruding.

SMovie 3 Krt4-GFP-KRas^V12^ cell basally extruding.

SMovie 4 Rotating view of projection of a KRT4-GFP-KRas^V12^ cell internalized between two layers of P63-positive basal cell layers.

SMovie 5 The nucleus of Krt4-GFP-KRas^V12^ cell that basally extruded gradually becomes more diffuse as it migrates in a H2afva:H2afva-mCherry zebrafish line.

SMovie 6 The nucleus of a migrating Krt4-GFP-KRas^V12^ cell in a H2afva:H2afva-mCherry zebrafish line has very weak Histone H2A signal.

SMovie 7 A GFP-KRas^V12^ cell pauses migrating, divides, and the daughter cells migrate away from each other.

SMovie 8 A GFP-KRas^V12^ cell sends out bipolar projections, similar to a neuron.

SMovie 9 A kdrl:mCherry zebrafish shows several GFP-KRas^V12^ cells within the blood vessels.

SMovie 10 Zoom of a movie showing a GFP-KRas^V12^ cell stuck within the blood vessel of a A kdrl:mCherry zebrafish.

SMovie 11 Zoom of a movie showing a GFP-KRas^V12^ impedes blood flow in a gata1:mCherry zebrafish.

SMovie 12 Extrusion of a GFP-KRas^V12^ ionocyte cell shows that it splits into an apical green portion and a red portion that migrates away.

SMovie 13 Extrusion of a GFP-KRas^V12^ periderm (outer epidermal) cell shows that it splits into an apical green portion and a red portion that migrates away.

SMovie 14 Extrusion of a krt4-GFP-KRas^V12^ epidermal cell shows that the apex fragments and the basal portion of the cell migrates away using a stable bleb with actin focused at the back.

SMovie 15 Unusual (only one found) of GFP-KRas^V12^ cells migrating away from an internalized mass.

**S Figure 1.**
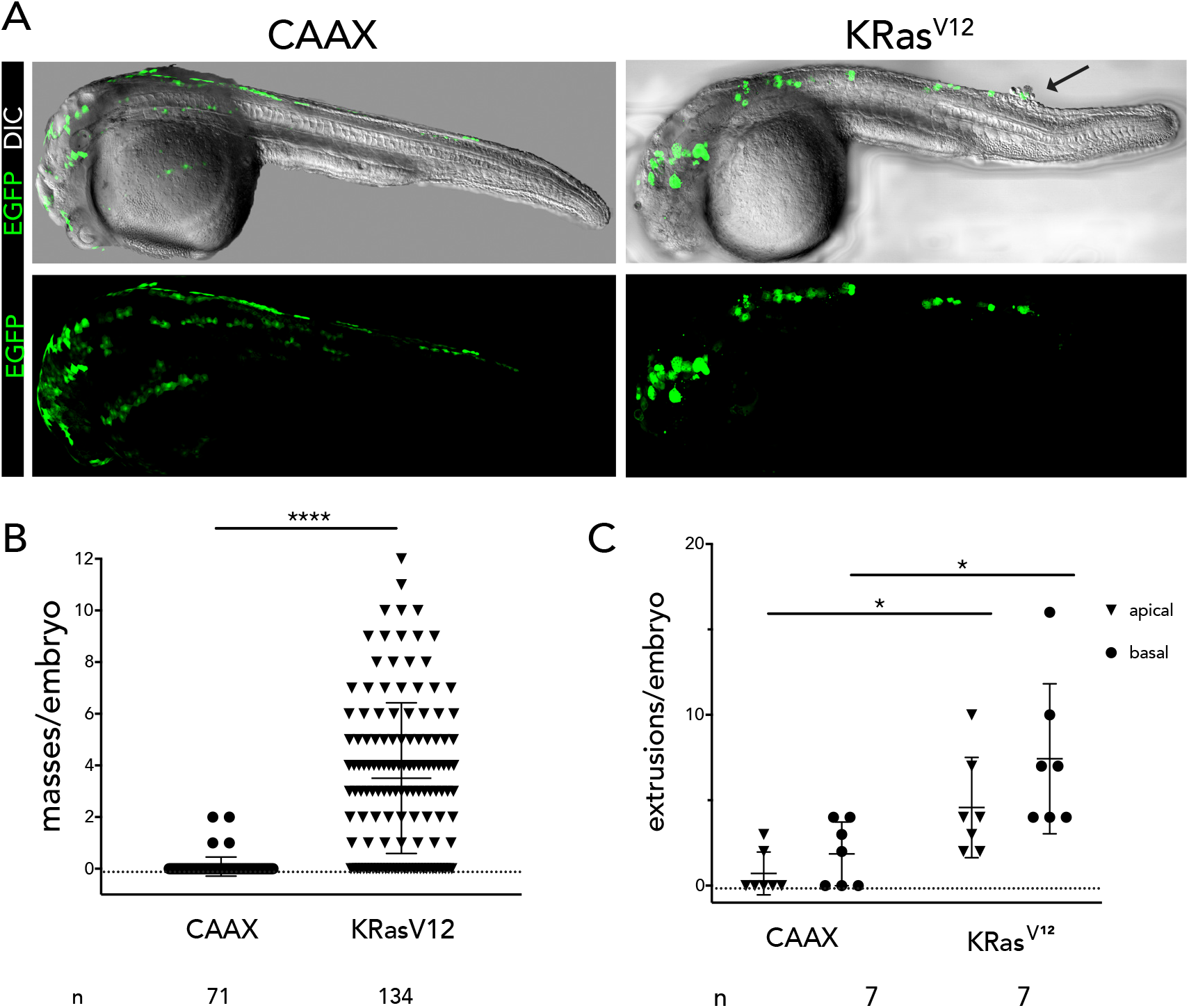
Loss of p53 enhances cell mass formation and extrusion rates of KRas^V12^ cells (A) 24 hpf p53^-/-^ embryos harboring EGFP-CAAX or KRas^V12^, where an epidermal cell mass is indicated by a black arrow. (B) Quantification of cell masses per embryo in p53^-/-^fish. (C) Quantification of live apical or basal extrusions per embryo in p53^-/-^fish. For all graphs, values are from the Ns listed below the graph, where error bars = s.e.m. P-values from unpaired T-tests compared to control are ^****^<0.0001, ^*^<0.01.

**SFigure 2.**
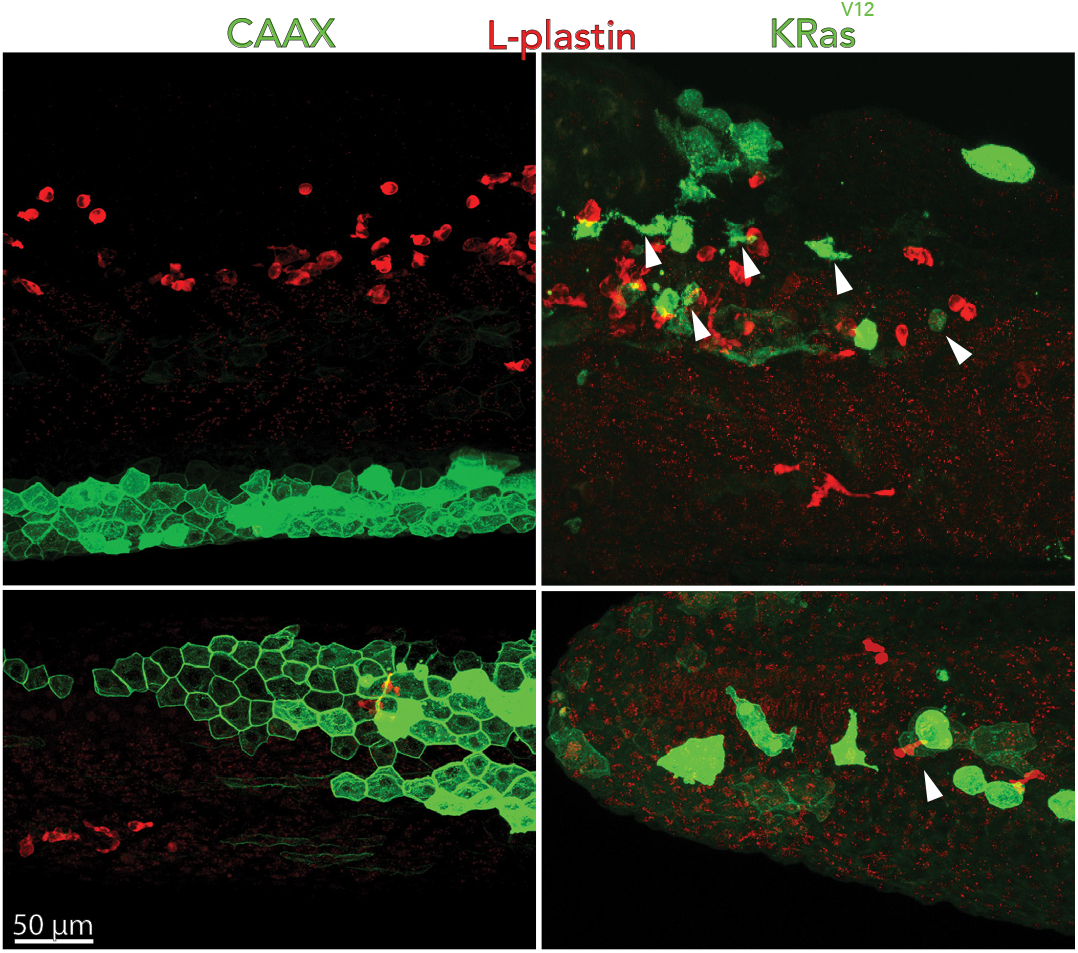
Macrophages do not colocalize with internalized, migrating KRas^V12^ cells Confocal projections showing that internalized EGFP-CAAX (A) or EGFP-KRas^V12^ (B) cells do not colocalize with macrophages (l-plastin, red) cells. While some of the cells appear to touch each other, they are not entirely engulfed by macrophages, indicating that their movement is not due to being engulfed by macrophages. Of 17 internalized cells quantified only one was ingested by a macrophage, consistent with the rate of dead internalized EGFP-KRas^V12^ in p53 mutant backgrounds.

## REFERENCES

Adams C.L., Chen Y.-T., Smith S.J., and James Nelson W. (1998). Mechanisms of Epithelial Cell–Cell Adhesion and Cell Compaction Revealed by High-resolution Tracking of E-Cadherin– Green Fluorescent Protein. The Journal of Cell Biology 142, 1105–1119.

Belaadi N., Aureille J., and Guilluy C. (2016). Under Pressure: Mechanical Stress Management in the Nucleus. Cells 5.

Boone B.A., Zeh H.J., 3rd, and Bahary N. (2018). Autophagy Inhibition in Pancreatic Adenocarcinoma. Clin Colorectal Cancer 17, 25–31.

Das S., and Batra S.K. (2015). Pancreatic cancer metastasis: are we being pre-EMTed? Curr Pharm Des 21, 1249–1255.

Demir I.E., Friess H., and Ceyhan G.O. (2015). Neural plasticity in pancreatitis and pancreatic cancer. Nat Rev Gastroenterol Hepatol 12, 649–659.

Eisenhoffer G.T., Loftus P.D., Yoshigi M., Otsuna H., Chien C.B., Morcos P.A., and Rosenblatt J. (2012). Crowding induces live cell extrusion to maintain homeostatic cell numbers in epithelia. Nature 484, 546–549.

Eisenhoffer G.T., and Rosenblatt J. (2011). Live imaging of cell extrusion from the epidermis of developing zebrafish. J Vis Exp.

Eisenhoffer G.T., and Rosenblatt J. (2013). Bringing balance by force: live cell extrusion controls epithelial cell numbers. Trends Cell Biol 23, 185–192.

Eisenhoffer G.T., Slattum G., Ruiz O.E., Otsuna H., Bryan C.D., Lopez J., Wagner D.S., Bonkowsky J.L., Chien C.B., Dorsky R.I., et al. (2017). A toolbox to study epidermal cell types in zebrafish. J Cell Sci 130, 269–277.

Elosegui-Artola A., Andreu I., Beedle, A.E.M., Lezamiz A., Uroz M., Kosmalska A.J., Oria R., Kechagia J.Z., Rico-Lastres P., Le Roux A.L., et al. (2017). Force Triggers YAP Nuclear Entry by Regulating Transport across Nuclear Pores. Cell 171, 1397–1410 e1314.

Fadul J., and Rosenblatt J. (2018). The forces and fates of extruding cells. Curr Opin Cell Biol 54, 66–71.

Fischer K.R., Durrans A., Lee S., Sheng J., Li F., Wong S.T., Choi H., El Rayes T., Ryu S., Troeger J., et al. (2015). Epithelial-to-mesenchymal transition is not required for lung metastasis but contributes to chemoresistance. Nature 527, 472–476.

Fouquet S., Lugo-Martinez V.H., Faussat A.M., Renaud F., Cardot P., Chambaz J., Pincon-Raymond M., and Thenet S. (2004). Early loss of E-cadherin from cell-cell contacts is involved in the onset of Anoikis in enterocytes. J Biol Chem 279, 43061–43069.

Frisch S.M., Schaller M., and Cieply B. (2013). Mechanisms that link the oncogenic epithelial-mesenchymal transition to suppression of anoikis. J Cell Sci 126, 21–29.

Gabriel M.T., Calleja L.R., Chalopin A., Ory B., and Heymann D. (2016). Circulating Tumor Cells: A Review of Non-EpCAM-Based Approaches for Cell Enrichment and Isolation. Clin Chem 62, 571–581.

Gu Y., Forostyan T., Sabbadini R., and Rosenblatt J. (2011). Epithelial cell extrusion requires the sphin-gosine-1-phosphate receptor 2 pathway. J Cell Biol 193, 667–676.

Gu Y., Shea J., Slattum G., Firpo M.A., Alexander M., Mulvihill S.J., Golubovskaya V.M., and Rosenblatt J. (2015). Defective apical extrusion signaling contributes to aggressive tumor hallmarks. eLife 4, e04069.

Gudipaty S.A., Lindblom J., Loftus P.D., Redd M.J., Edes K., Davey C.F., Krishnegowda V., and Rosenblatt J. (2017). Mechanical stretch triggers rapid epithelial cell division through Piezo1. Nature 543, 118–121.

Gudipaty S.A., and Rosenblatt J. (2017). Epithelial cell extrusion: Pathways and pathologies. Semin Cell Dev Biol 67, 132–140.

Hanahan D., and Weinberg R.A. (2011). Hallmarks of cancer: the next generation. Cell 144, 646–674.

Hartenstein V., Younossi-Hartenstein A., and Lekven A. (1994). Delamination and division in the Drosophila neurectoderm: spatiotemporal pattern, cytoskeletal dynamics, and common control by neurogenic and segment polarity genes. Dev Biol 165, 480–499.

Hendley A.M., Wang Y.J., Polireddy K., Alsina J., Ahmed I., Lafaro K.J., Zhang H., Roy N., Savidge S.G., Cao Y., et al. (2016). p120 Catenin Suppresses Basal Epithelial Cell Extrusion in Invasive Pancreatic Neoplasia. Cancer Res 76, 3351–3363.

Hingorani S.R., Wang L., Multani A.S., Combs C., Deramaudt T.B., Hruban R.H., Rustgi A.K., Chang S., and Tuveson D.A. (2005). Trp53R172H and KrasG12D cooperate to promote chromosomal instability and widely metastatic pancreatic ductal adenocarcinoma in mice. Cancer Cell 7, 469–483.

Huisken J., and Stainier D.Y. (2007). Even fluorescence excitation by multidirectional selective plane illumination microscopy (mSPIM). Opt Lett 32, 2608–2610.

Ivanovska I.L., Shin J.W., Swift J., and Discher D.E. (2015). Stem cell mechanobiology: diverse lessons from bone marrow. Trends Cell Biol 25, 523–532.

Kalluri R., and Weinberg R.A. (2009). The basics of epithelial-mesenchymal transition. J Clin Invest 119, 1420–1428.

Kang R., and Tang D. (2012). Autophagy in pancreatic cancer pathogenesis and treatment. Am J Cancer Res 2, 383–396.

Kaufmann A., Mickoleit M., Weber M., and Huisken J. (2012). Multilayer mounting enables long-term imaging of zebrafish development in a light sheet microscope. Development 139, 3242–3247.

Kumar S., Park S.H., Cieply B., Schupp J., Killiam E., Zhang F., Rimm D.L., and Frisch S.M. (2011). A pathway for the control of anoikis sensitivity by E-cadherin and epithelial-to-mesenchymal transition. Mol Cell Biol 31, 4036–4051.

Liu Y.J., Le Berre M., Lautenschlaeger F., Maiuri P., Callan-Jones A., Heuze M., Takaki T., Voituriez R., and Piel M. (2015a). Confinement and low adhesion induce fast amoeboid migration of slow mesenchymal cells. Cell 160, 659–672.

Liu Y.J., Le Berre M., Lautenschlaeger F., Maiuri P., Callan-Jones A., Heuze M., Takaki T., Voituriez R., and Piel M. (2015b). Confinement and low adhesion induce fast amoeboid migration of slow mesenchymal cells. Cell 160, 659–672.

Marshall T.W., Lloyd I.E., Delalande J.M., Nathke I., and Rosenblatt J. (2011). The tumor suppressor adenomatous polyposis coli controls the direction in which a cell extrudes from an epithelium. Mol Biol Cell 22, 3962–3970.

McGregor A.L., Hsia C.R., and Lammerding J. (2016). Squish and squeeze-the nucleus as a physical barrier during migration in confined environments. Curr Opin Cell Biol 40, 32–40.

Meyer A.S., Miller M.A., Gertler F.B., and Lauffenburger D.A. (2013). The receptor AXL diversifies EGFR signaling and limits the response to EGFR-targeted inhibitors in triple-negative breast cancer cells. Sci Signal 6, ra66.

Moore P.S., Beghelli S., Zamboni G., and Scarpa A. (2003). Genetic abnormalities in pancreatic cancer. Mol Cancer 2, 7.

Mosimann C., Puller A.C., Lawson K.L., Tschopp P., Amsterdam A., and Zon L.I. (2013). Site-directed zebrafish transgenesis into single landing sites with the phiC31 integrase system. Dev Dyn 242, 949–963.

Muthuswamy S.K., and Xue B. (2012). Cell polarity as a regulator of cancer cell behavior plasticity. Annu Rev Cell Dev Biol 28, 599–625.

Neben K., Hübner G., Folprecht G., Jäger D., and Krämer A. (2008). Metastases in the Absence of a Primary Tumor: Advances in the Diagnosis and Treatment of CUP Syndrome. Deutsches Ärzteblatt International 105, 733–740.

Rasheed Z.A., Matsui W., and Maitra A. (2012). Pathology of pancreatic stroma in PDAC. In Pancreatic Cancer and Tumor Microenvironment P.J. Grippo, and H.G. Munshi, eds. (Trivandrum (India)).

Rhim A.D., Mirek E.T., Aiello N.M., Maitra A., Bailey J.M., McAllister F., Reichert M., Beatty G.L., Rustgi A.K., Vonderheide R.H., et al. (2012). EMT and Dissemination Precede Pancreatic Tumor Formation. Cell 148, 349–361.

Robu M.E., Larson J.D., Nasevicius A., Beiraghi S., Brenner C., Farber S.A., and Ekker S.C. (2007). p53 activation by knockdown technologies. PLoS Genet 3, e78.

Rosenblatt J., Raff M.C., and Cramer L.P. (2001). An epithelial cell destined for apoptosis signals its neighbors to extrude it by an actin- and myosin-dependent mechanism. Curr Biol 11, 1847–1857.

Ruprecht V., Wieser S., Callan-Jones A., Smutny M., Morita H., Sako K., Barone V., Ritsch-Marte M., Sixt M., Voituriez R., et al. (2015a). Cortical contractility triggers a stochastic switch to fast amoeboid cell motility. Cell 160, 673–685.

Ruprecht V., Wieser S., Callan-Jones A., Smutny M., Morita H., Sako K., Barone V., Ritsch-Marte M., Sixt M., Voituriez R., et al. (2015b). Cortical contractility triggers a stochastic switch to fast amoeboid cell motility. Cell 160, 673–685.

Saito Y., Desai R.R., and Muthuswamy S.K. (2018). Reinterpreting polarity and cancer: The changing landscape from tumor suppression to tumor promotion. Biochim Biophys Acta 1869, 103–116.

Saloman J.L., Albers K.M., Li D., Hartman D.J., Crawford H.C., Muha E.A., Rhim A.D., and Davis B.M. (2016). Ablation of sensory neurons in a genetic model of pancreatic ductal adenocarcinoma slows initiation and progression of cancer. Proc Natl Acad Sci U S A 113, 3078–3083.

Scarpa A., Capelli P., Mukai K., Zamboni G., Oda T., Iacono C., and Hirohashi S. (1993). Pancreatic adenocarcinomas frequently show p53 gene mutations. Am J Pathol 142, 1534–1543.

Scherz-Shouval R., Weidberg H., Gonen C., Wilder S., Elazar Z., and Oren M. (2010). p53-dependent regulation of autophagy protein LC3 supports cancer cell survival under prolonged starvation. Proc Natl Acad Sci U S A 107, 18511–18516.

Schindelin J., Arganda-Carreras I., Frise E., Kaynig V., Longair M., Pietzsch T., Preibisch S., Rueden C., Saalfeld S., Schmid B., et al. (2012). Fiji: an open-source platform for biological-image analysis. Nature Methods 9, 676.

Schmick M., Vartak N., Papke B., Kovacevic M., Truxius D.C., Rossmannek L., and Bastiaens, P.I.H. (2014). KRas localizes to the plasma membrane by spatial cycles of solubilization, trapping and vesicular transport. Cell 157, 459–471.

Schmid B., Shah G., Scherf N., Weber M., Thierbach K., Campos C.P., Roeder I., Aanstad P., and Huisken J. (2013). High-speed panoramic light-sheet microscopy reveals global endodermal cell dynamics. Nat Commun 4, 2207.

Simoes S., Oh Y., Wang, M.F.Z., Fernandez-Gonzalez R., and Tepass U. (2017). Myosin II promotes the anisotropic loss of the apical domain during Drosophila neuroblast ingression. J Cell Biol 216, 1387–1404.

Slattum G., Gu Y., Sabbadini R., and Rosenblatt J. (2014). Autophagy in oncogenic K-Ras promotes basal extrusion of epithelial cells by degrading S1P. Curr Biol 24, 19–28.

Slattum G., McGee K.M., and Rosenblatt J. (2009). P115 RhoGEF and microtubules decide the direction apoptotic cells extrude from an epithelium. J Cell Biol 186, 693–702.

Slattum G.M., and Rosenblatt J. (2014). Tumour cell invasion: an emerging role for basal epithelial cell extrusion. Nat Rev Cancer 14, 495–501.

Swift J., Ivanovska I.L., Buxboim A., Harada T., Dingal P.C., Pinter J., Pajerowski J.D., Spinler K.R., Shin J.W., Tewari M., et al. (2013). Nuclear lamin-A scales with tissue stiffness and enhances matrix-directed differentiation. Science 341, 1240104.

Weber M., Mickoleit M., and Huisken J. (2014). Multilayer mounting for long-term light sheet microscopy of zebrafish. J Vis Exp, e51119.

Welch M.D. (2015). Cell migration, freshly squeezed. Cell 160, 581–582.

Wilson J.L., Kefaloyianni E., Stopfer L., Harrison C., Sabbisetti V.S., Fraenkel E., Lauffenburger D.A., and Herrlich A. (2018). Functional Genomics Approach Identifies Novel Signaling Regulators of TGFalpha Ecto-domain Shedding. Mol Cancer Res 16, 147–161.

Ye X., and Weinberg R.A. (2015). Epithelial-Mesenchymal Plasticity: A Central Regulator of Cancer Progression. Trends Cell Biol 25, 675–686.

Zheng X., Carstens J.L., Kim J., Scheible M., Kaye J., Sugimoto H., Wu C.C., LeBleu V.S., and Kalluri R. (2015). Epithelial-to-mesenchymal transition is dispensable for metastasis but induces chemoresistance in pancreatic cancer. Nature 527, 525–530.

